## Supplemental Figures and Tables for "A putative cap binding protein and the methyl phosphate capping enzyme Bin3/MePCE function in telomerase biogenesis"

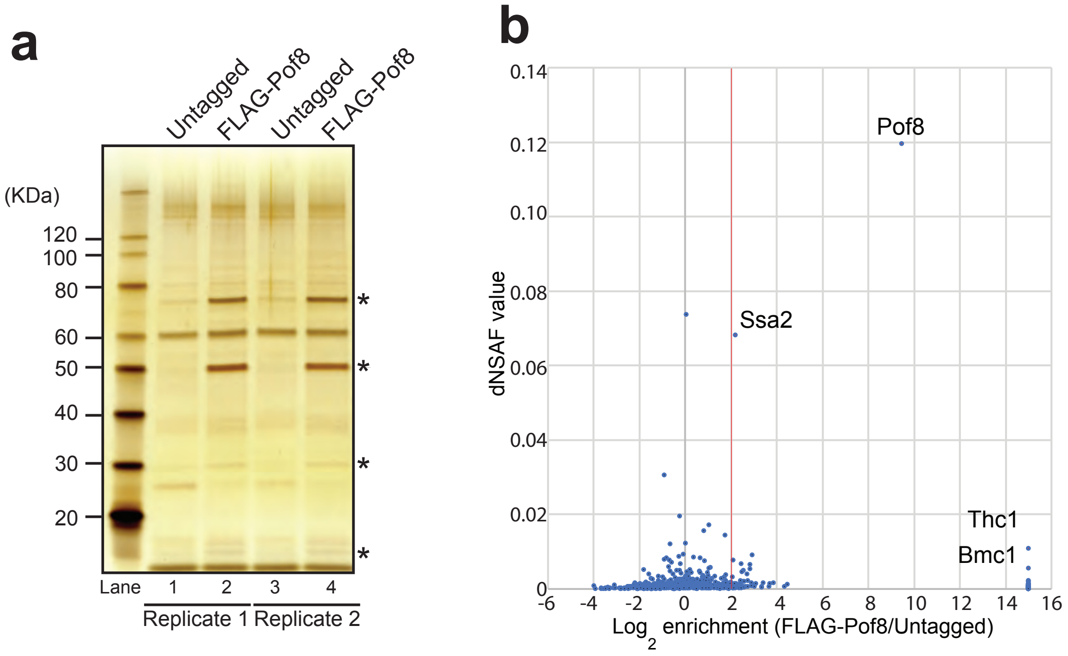


**Supplementary Figure 1.** Identification of Pof8 interacting proteins. **a** Silver stained SDS-PAGE of immunoprecipitations (IPs) for 3xFLAG-Pof8 expressed from a plasmid under the control of its endogenous promoter. The two bands marked with asterisks at ~50 kDa and ~75 kDa represent 3xFLAG-Pof8 and Ssa2, a heat shock protein, respectively. Two biological replicates are shown. Two lower bands are more prominent in lanes 2 and 4 compared to lanes 1 and 3 and are also marked with asterisks. **b** Scatter plot of proteins enriched in 3xFLAG-Pof8 IP by mass spectrometry. The x-axis shows the average enrichment in the 3xFLAG-Pof8 samples compared to untagged controls. Proteins not detected in the controls have a calculated enrichment value of infinity. To plot these data points, the enrichment value was arbitrarily set to $2^{15}$. On the y-axis is the distributed Normalized Spectral Abundance Factor (dNSAF) mean value from tagged samples. The vertical red line marks a 4-fold enrichment in the tagged over untagged samples.

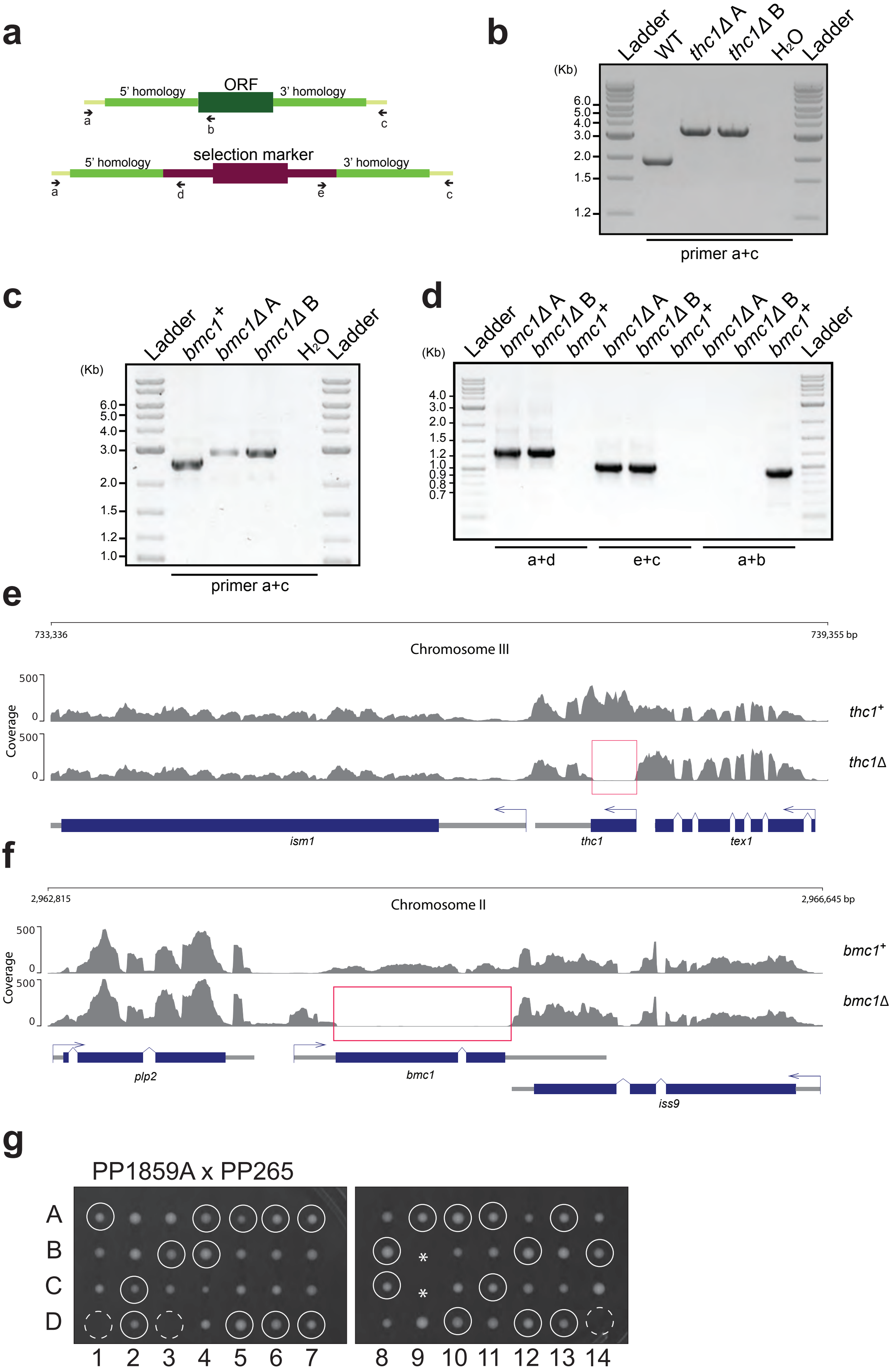
a
5' homology
ORF
3' homology
a
b
c
selection marker
d
e
b
Ladder
WT
thc1Δ A
thc1Δ B
H2O
(Kb)
6.0
5.0
4.0
3.0
2.0
1.5
1.2
primer a+c
c
bmc1+
bmc1Δ A
bmc1Δ B
1.0
d
a+d
e+c
a+b
0.9
0.8
0.7
e
733,336
739,355 bp
Chromosome III
500
0
Coverage
thc1+
thc1Δ
ism1
thc1
tex1
f
2,962,815
2,966,645 bp
Chromosome II
bmc1+
bmc1Δ
plp2
bmc1
iss9
g
PP1859A x PP265
A
B
C
D
1 2 3 4 5 6 7
8 9 10 11 12 13 14

**Supplementary Figure 2. a** Schematic of primers used to verify the deletion of *thc1* and *bmc1*, respectively. **b** Diagnostic PCR across the *thc1* locus using genomic DNA from two *thc1D* isolates and a *thc1*$^{+}$ control strain and primers a and c as indicated in the schematic in a. The lane labelled $H_2O$ contained no genomic DNA. **c** Diagnostic PCR as in (b) but for the *bmc1* locus. **d** Diagnostic PCR to confirm the insertion of the *bmc1* knock-out cassette at the correct location in the genome. Primers a+d give a product of the expected size if insertion of the 5' homology occurred in the correct location; primers e+c give a product of the expected size if insertion of the 5' homology occurred in the correct location. Primers a+b only give a product if the *bmc1* open reading frame is present. **e** Coverage tracks from RNA-sequencing analysis for the *thc1* locus in a *thc1Δ* and *thc1*$^{+}$ control strain. The absence of reads mapping to the *thc1* open reading frame in the deletion strain boxed in red serves as an independent confirmation of the locus having been successfully deleted. **f** Coverage tracks from RNA sequencing analysis for the *bmc1* locus in a *bmc1Δ* and *bmc1*$^{+}$ control strain. **g** Tetrad dissection of a cross between *bmc1Δ* strain PP1859 and PP265 (wildtype *S. pombe* Lindner 972). Each column represents the spores from one tetrad. White circles denote *bmc1Δ* colonies based on growth following replica-plating to YEA NAT media. Dashed circles indicate *bmc1Δ* colonies that did not grow, the two asterisks in tetrad 9 label the positions of a *bmc1*$^{+}$ and a *bmc1Δ* spore that did not form colonies.

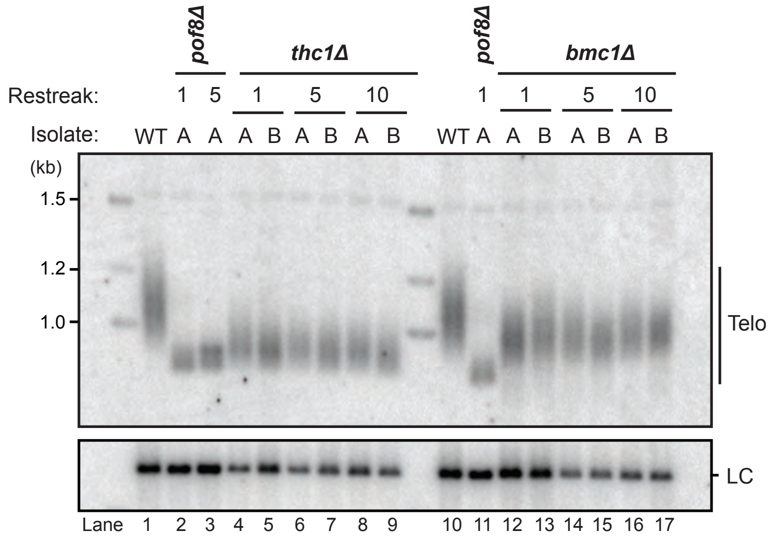


**Supplementary Figure 3.** Stable but short telomeres in thc1 and bmc1 deletion strains. Telomeric Southern blot comparing telomere length in wildtype (WT) and *thc1*Δ or *bmc1*Δ cells. Two independent isolates of each deletion (A and B) were restreaked ten times in series and telomere length was analyzed (one restreak equals 20-25 generations). A *pof8*Δ strain was included as control. The *rad16*$^{+}$ locus was probed to assess integrity and equal loading of the genomic DNA samples.

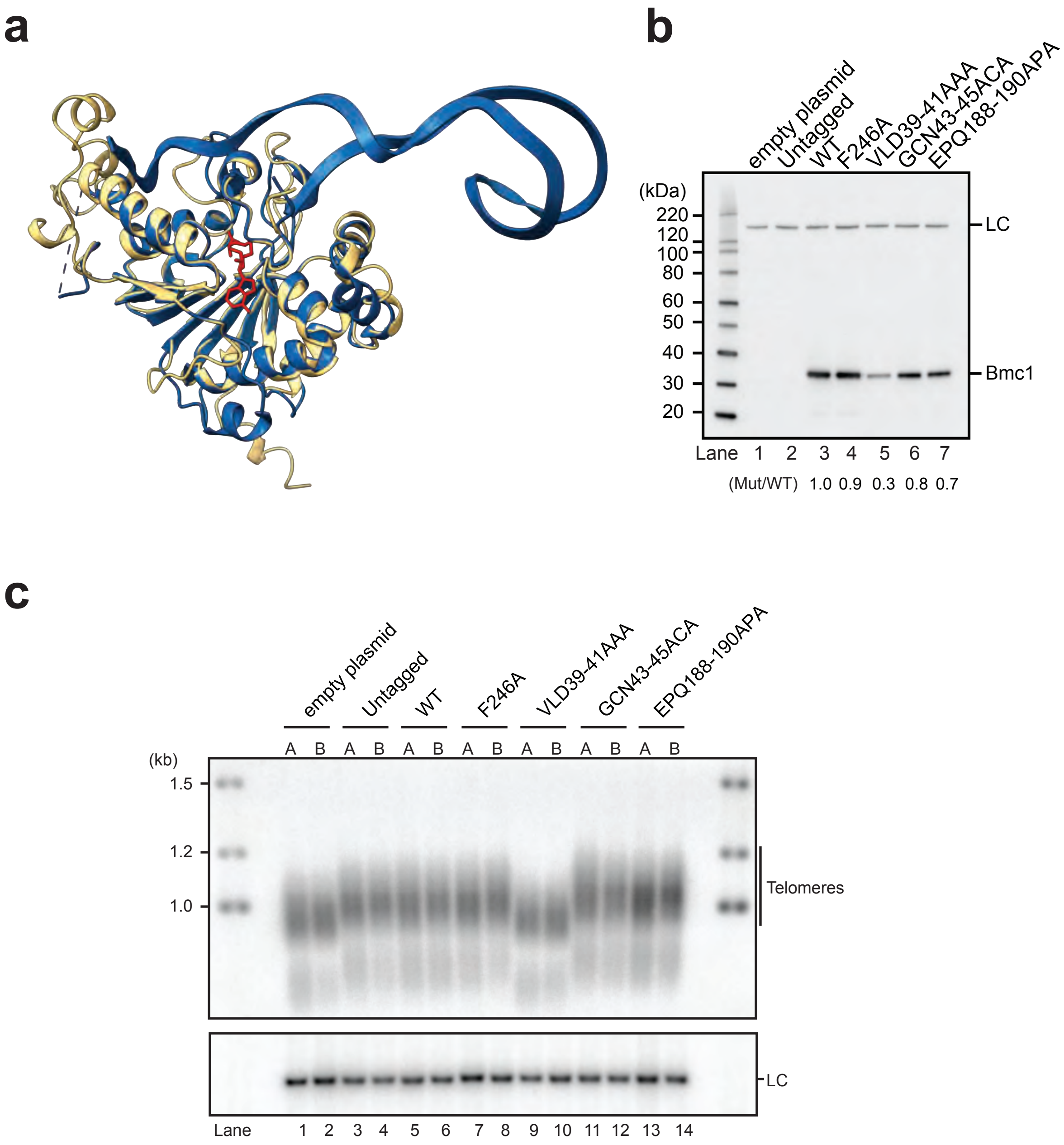


**Supplementary Figure 4.** The function of Bmc1 in telomerase biogenesis is independent of its catalytic activity. **a** ChimeraX-1.25 overlay of the Alphafold predicted structure of *S. pombe* Bmc1 (orange) and the human MePCE methyltransferase domain bound to S-adenosylhomocysteine (blue, PDB ID 6DCB). S-adenosylhomocysteine shown in red. **b** Western blot analysis of Twinstrep-tagged wildtype (WT) and mutant Bmc1 designed to affect the methyltransferase activity using α-Strep-tag II antibody. All versions of Bmc1 were expressed from plasmids under the control of the endogenous promoter in a *bmc1*Δ background. A non-specific band recognized by α-Strep-tag II in the absence of the epitope-tag (lane 2) was used as intrinsic loading control (LC). Quantification of mutants compared with WT are shown below the lane numbers. **c** Telomeric Southern blot for *bmc1* mutants. Two independent isolates of each strain are shown. A probe against the *rad16*$^{+}$ locus was used as LC. Lane numbers are indicated below the blot.

### Paez et al.; Supplementary Figure 5

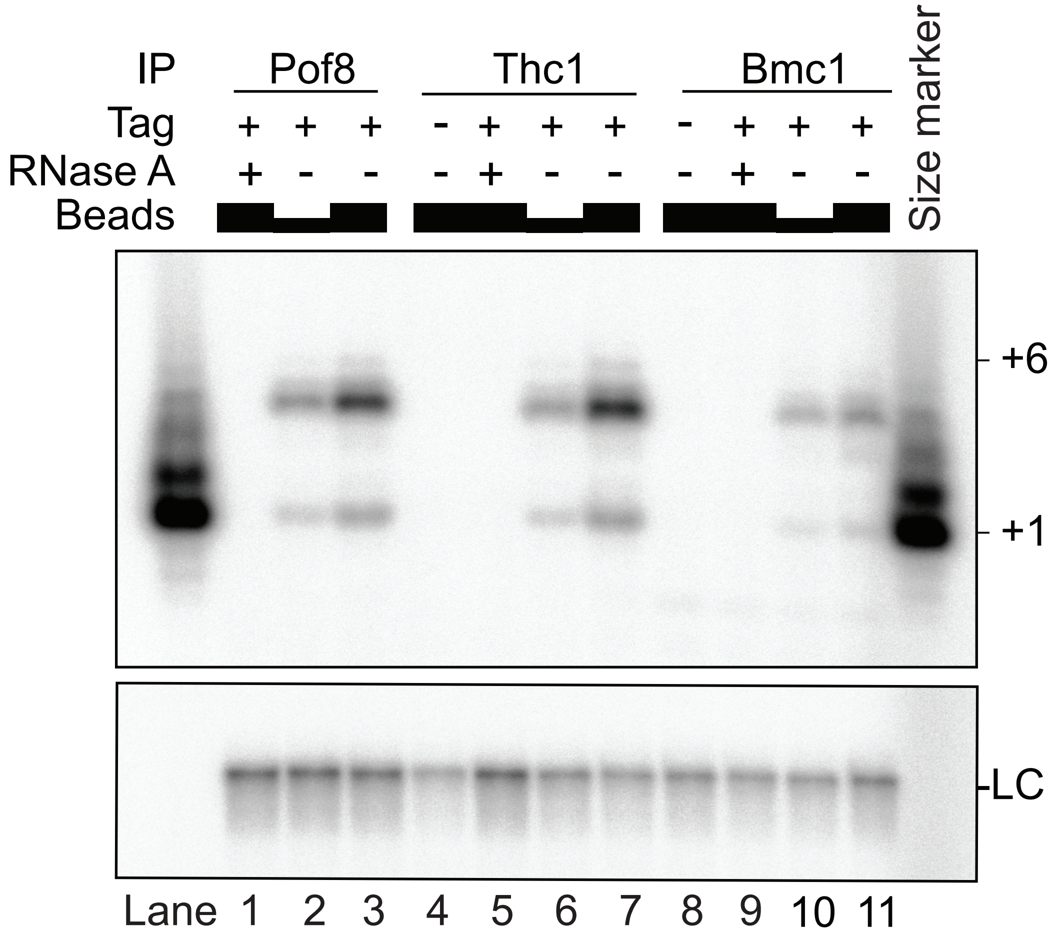


**Supplementary Figure 5.** Thc1 and Bmc1 are associated with telomerase activity. Activity assays were performed following immunoprecipation of 3xFLAG-Pof8 with anti-FLAG antibody coated Dynabeads, Thc1-2xV5 with anti-V5 antibody coated Dynabeads and Bmc1-Twinstrep with StrepTactin Sepharose. 10 µL (lanes 2, 6 and 10) or 20 µL (lanes 1, 3-5, 7-9 and 11) of IP suspensions were used for telomerase activity assay. A 32P labelled 100-mer oligo nucleotide was used as precipitation and loading control (LC).

#### Paez et al.; Supplementary Figure 6

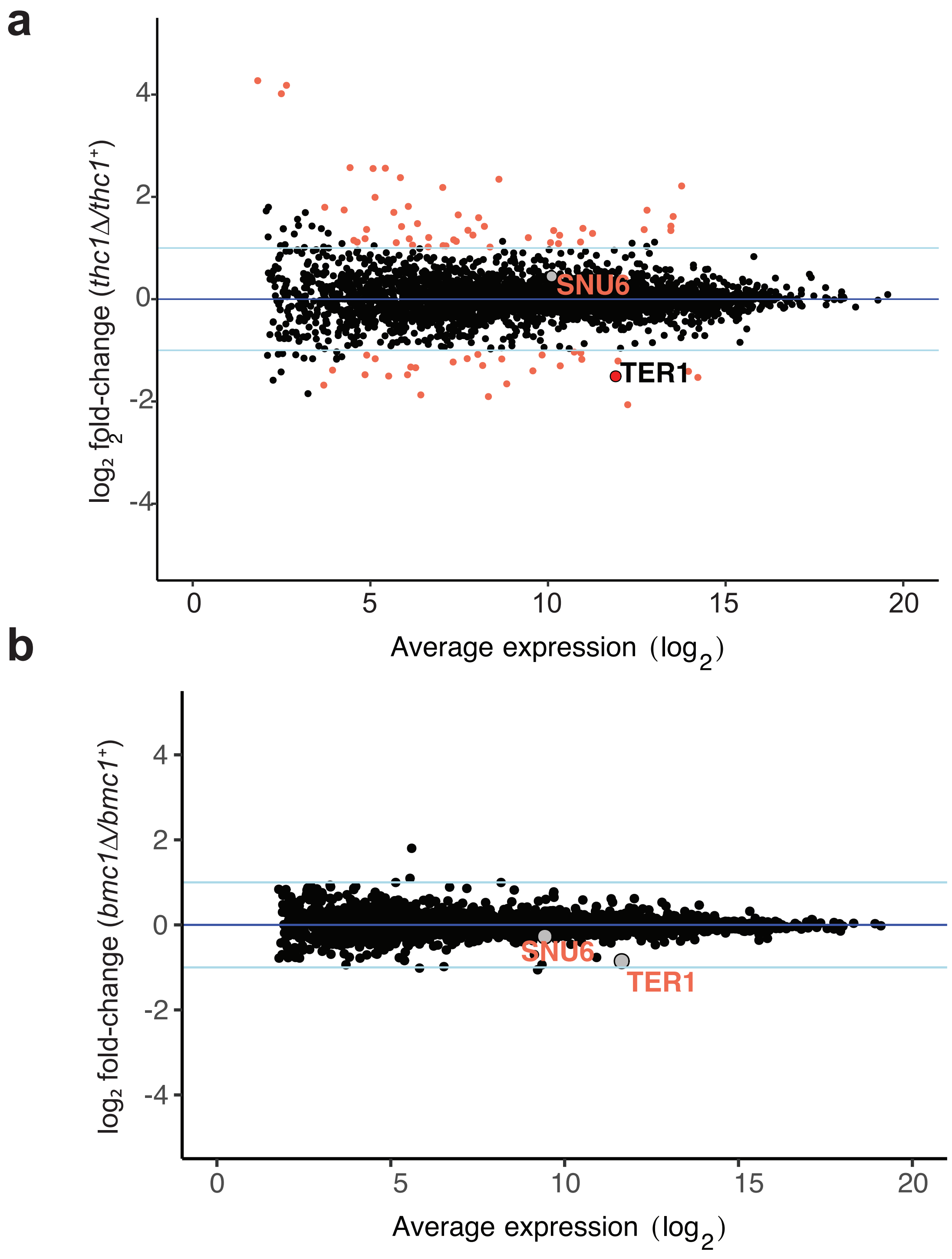


**Supplementary Figure 6.** MA plot of differential expression analysis for the deletion of *thc1* (**a**) and *bmc1* (**b**). Average expression is plotted on the x-axis and $\log_2$ fold change between the deletion and wildtype is plotted on the y-axis. Differentially expressed genes with an absolute $\log_2$ fold change of ≥ 1 and an adjusted p-value < 0.05 are colored in orange.

**a**

Figure 2a Figure 2b Figure 2c

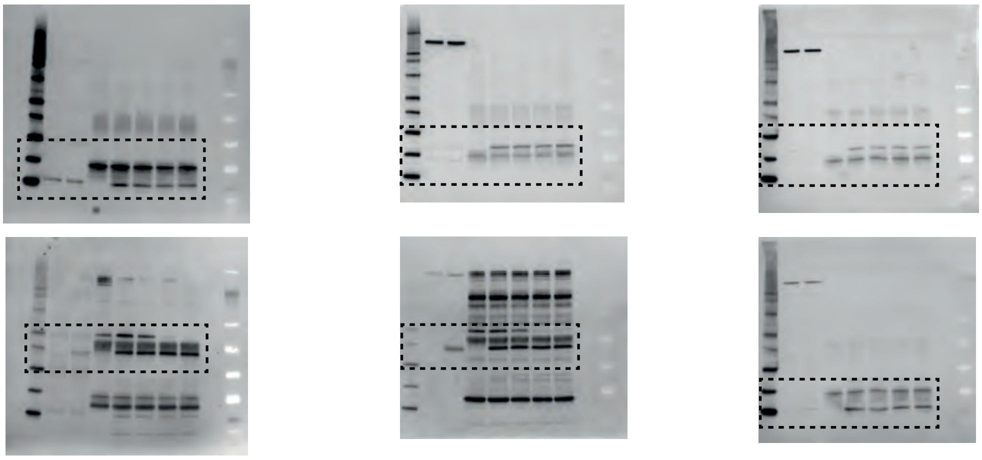

**b**

Figure 3d

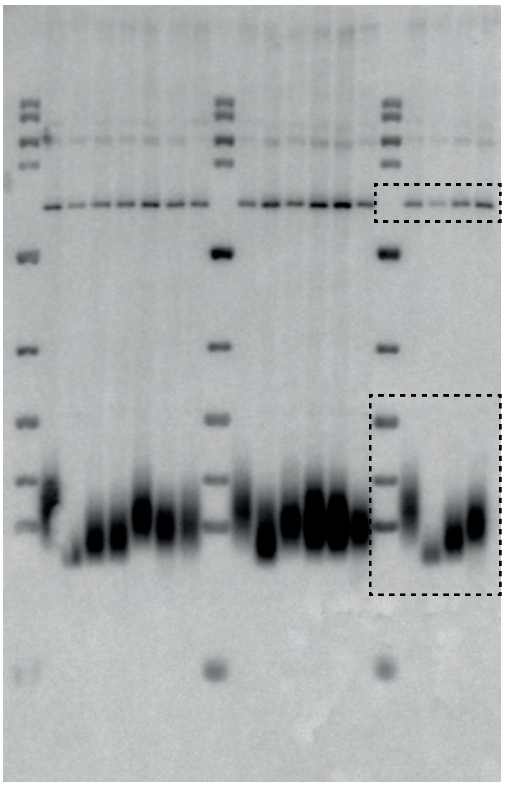

**c**

Figure 4b

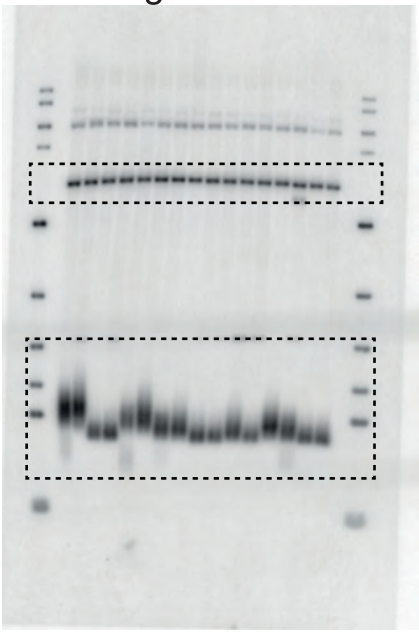

Figure 4c

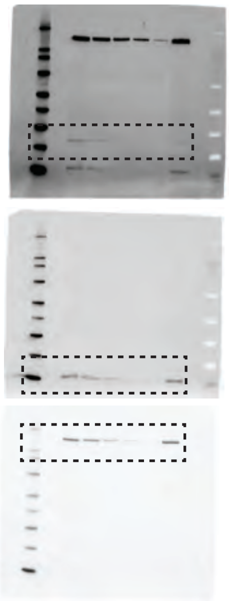

Figure 4d

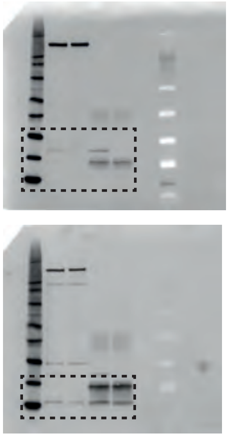

Figure 4e

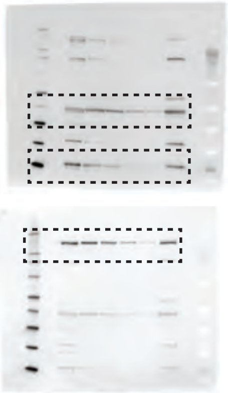

Figure 4f

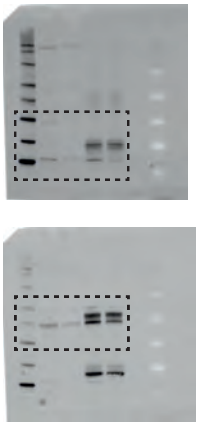

Figure 4g

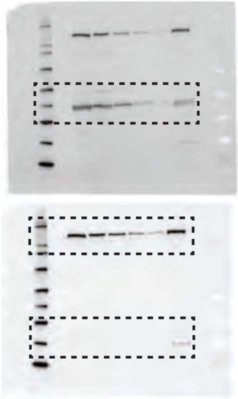

Figure 4h

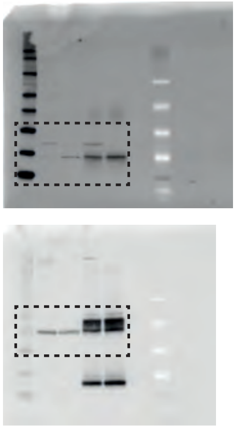

**d**

Figure 5a

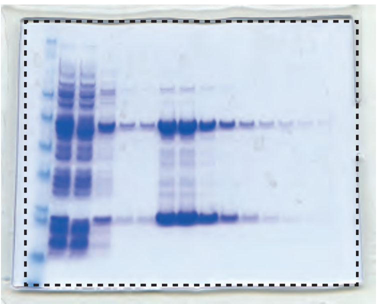

Figure 5b

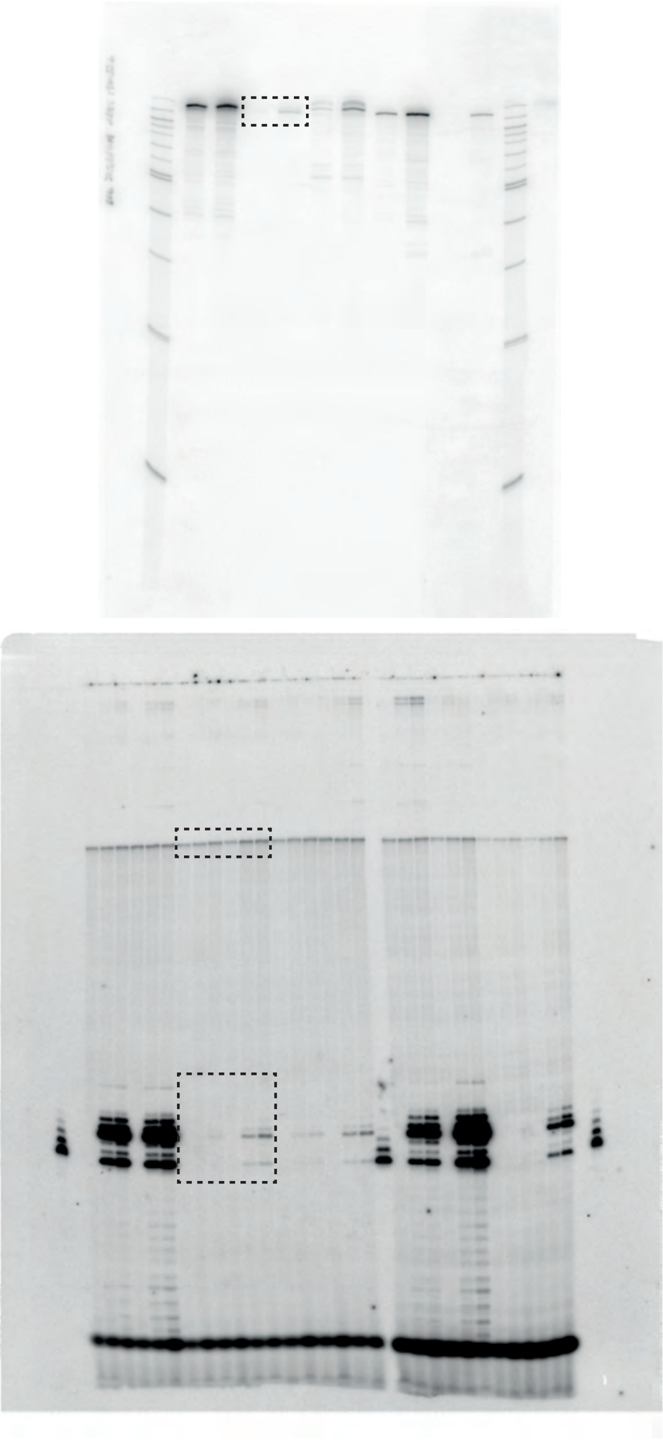

Figure 5c

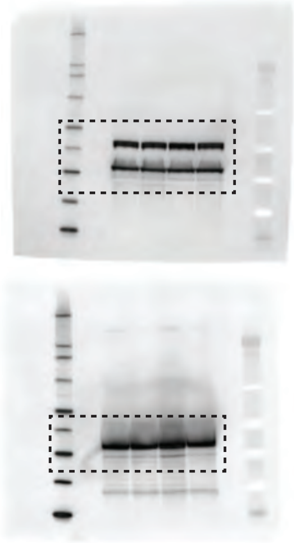

Figure 5e

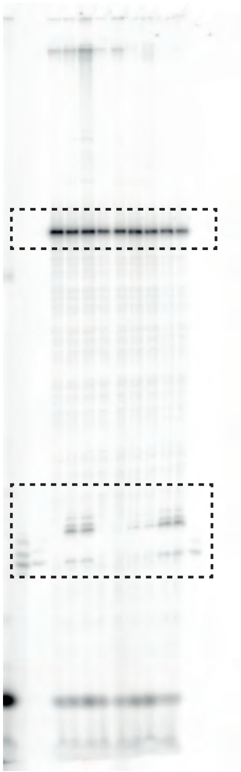

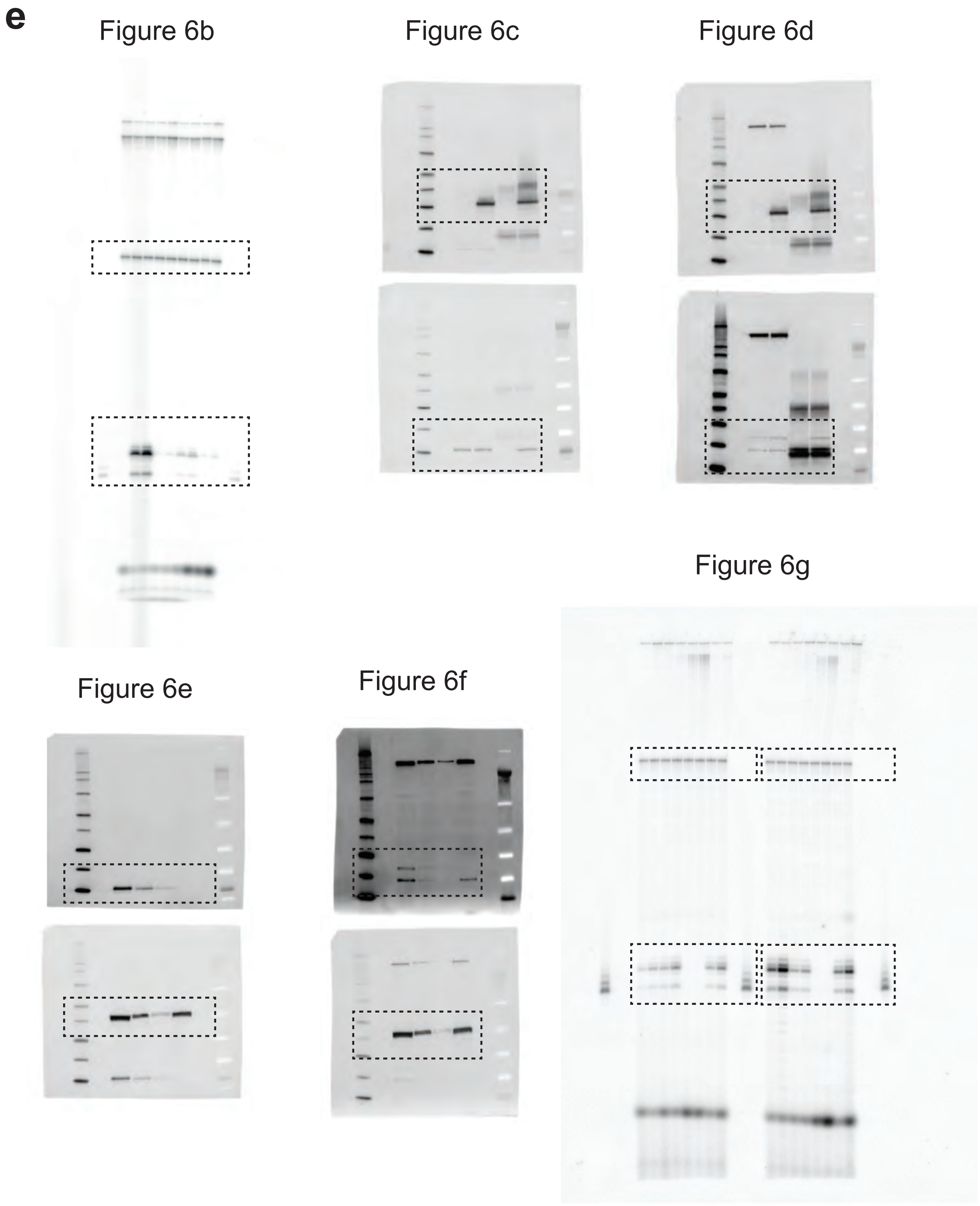


**Supplementary Figure 7.** Raw data presented in main figures. **a** Uncropped blots from Figure 2a-c. **b** Figure 3d. **c** Figure 4b-h. **d** Figure 5a-c, e. **e** Figure 6b-g.

**Supplementary Table 1:** Proteins enriched in Pof8-FLAG affinity purification with average dNSAF values >0.005 for the tagged Pof8 samples. Proteins are sorted by average fold enrichment in tagged Pof8 over untagged control.

| NCBI_Gene | gene_ID | Description | Pof8 avg dNSAF | control avg dNSAF | log2 fold change Pof8/ctrl | MW (Da) |
|---|---|---|---|---|---|---|
| | SPCC18B5.09c | sequence orphan | 0.010803 | 0 | ∞ (15) | 13507 |
| | SPBC2A9.10 | Bin3 family, transcriptional and translational regulator (predicted) | 0.005521 | 0 | ∞ (15) | 30485 |
| pof8 | SPAC17G6.17 | F-box protein Pof8 | 0.119549 | 0.000168 | 9.47 | 46808 |
| leu1 | SPBC1A4.02c | 3-isopropylmalate dehydrogenase Leu1 | 0.008942 | 0.001156 | 2.95 | 39733 |
| ubi5 | SPAC589.10c | ubiquitin-40S ribosomal protein S31 fusion protein | 0.006532 | 0.000904 | 2.85 | 17215 |
| rvb2 | SPBC83.08 | AAA family ATPase Rvb2 | 0.005249 | 0.000729 | 2.85 | 51562 |
| hsp90 | SPAC926.04c | Hsp90 chaperone | 0.006587 | 0.001186 | 2.47 | 80596 |
| ssa2 | SPCC1739.13 | heat shock protein Ssa2 (predicted) | 0.068151 | 0.014617 | 2.22 | 70233 |
| mcp60 | SPAC12G12.04 | mitochondrial heat shock protein Hsp60/Mcp60 | 0.007729 | 0.001784 | 2.12 | 62168 |
| rpp203 | SPAC1071.08 | 60S acidic ribosomal protein P2C | 0.014391 | 0.004215 | 1.77 | 11114 |
| rps3 | SPBC16G5.14c | 40S ribosomal protein S3 | 0.005661 | 0.002627 | 1.11 | 27553 |
| tdh1 | SPBC32F12.11 | glyceraldehyde-3-phosphate dehydrogenase Tdh1 | 0.005581 | 0.002685 | 1.06 | 35870 |
| rpp201 | SPBP8B7.06 | 60S acidic ribosomal protein P2A | 0.017156 | 0.008308 | 1.05 | 11158 |
| ssc1 | SPAC664.11 | mitochondrial heat shock protein Hsp70 | 0.00747 | 0.004123 | 0.86 | 72977 |
| sks2 | SPBC1709.05 | heat shock protein, ribosome associated molecular chaperone Sks2 | 0.015436 | 0.008584 | 0.85 | 67206 |
| | SPAC12G12.07c | conserved fungal protein | 0.008371 | 0.006479 | 0.37 | 45724 |
| lsm4 | SPBC30D10.06 | U6 snRNP-associated protein Lsm4 (predicted) | 0.005178 | 0.004473 | 0.21 | 13941 |
| cxr1 | SPBC23E6.01c | mRNA processing factor (predicted) | 0.07353 | 0.070509 | 0.06 | 51705 |
| pabp | SPAC57A7.04c | mRNA export shuttling protein | 0.012228 | 0.011967 | 0.03 | 71513 |

**Supplementary Table 2: Genes affected in expression level by the deletion of *thc1***

**a** Genes with RNA levels decreased by more than 2-fold in the absence of *thc1*$^+$ based on DESeq2 analysis of RNA samples from triplicate cultures of *thc1*$^+$ and *thc1Δ* cells. A two-sided Wald test was used and adjusted *p*-values were calculated by Benjamini-Hochberg **b** Genes with increased expression by more than 2-fold by the same analysis as in (a).

**a**

| gene_ID | external_gene_ID | log2_FC | adj. p-value | gene_biotype |
|---|---|---|---|---|
| SPNCRNA.863 | SPNCRNA.863 | -2.0640 | 9.6E-141 | ncRNA |
| SPNCRNA.1344 | SPBC1271.08c-antisense-1 | -1.9046 | 6.7E-34 | ncRNA |
| SPCC569.02c | SPCC569.02c | -1.8719 | 1.2E-12 | protein_coding |
| SPBCPT2R1.07c | SPBCPT2R1.07c | -1.6810 | 2.5E-02 | pseudogene |
| SPNCRNA.742 | SPAC9.08c-antisense-1 | -1.6559 | 3.5E-16 | ncRNA |
| SPBP4G3.02 | pho1 | -1.5304 | 2.7E-236 | protein_coding |
| SPNCRNA.214 | ter1 | -1.5083 | 7.8E-128 | ncRNA |
| SPNCRNA.923 | SPNCRNA.923 | -1.5039 | 1.5E-05 | ncRNA |
| SPNCRNA.130 | omt3 | -1.4787 | 4.7E-06 | ncRNA |
| SPNCRNA.1255 | SPNCRNA.1255 | -1.4776 | 6.9E-04 | ncRNA |
| SPAC821.10c | sod1 | -1.4115 | 1.0E-138 | protein_coding |
| SPCC70.08c | SPCC70.08c | -1.4017 | 1.2E-42 | protein_coding |
| SPCC663.14c | trp663 | -1.3878 | 4.1E-02 | protein_coding |
| SPCC1393.14 | ten1 | -1.3405 | 5.2E-07 | protein_coding |
| SPNCRNA.1605 | SPNCRNA.1605 | -1.3271 | 3.2E-07 | ncRNA |
| SPCC18B5.09c | thc1 | -1.3033 | 2.0E-76 | protein_coding |
| SPNCRNA.888 | end4-antisense-1 | -1.2969 | 8.5E-23 | ncRNA |
| SPAC186.06 | SPAC186.06 | -1.2292 | 5.1E-09 | protein_coding |
| SPNCRNA.1626 | SPNCRNA.1626 | -1.2100 | 4.9E-44 | ncRNA |
| SPBC1271.07c | SPBC1271.07c | -1.1767 | 4.9E-24 | protein_coding |
| SPBPB2B2.01 | SPBPB2B2.01 | -1.1699 | 1.4E-24 | protein_coding |
| SPBPJ4664.03 | mfm3 | -1.1656 | 4.3E-03 | protein_coding |
| SPNCRNA.1340 | SPNCRNA.1340 | -1.1631 | 5.2E-15 | ncRNA |
| SPAC8E11.12 | SPAC8E11.12 | -1.0941 | 1.5E-02 | protein_coding |
| SPAC27D7.03c | mei2 | -1.0916 | 3.0E-17 | protein_coding |
| SPNCRNA.944 | SPNCRNA.944 | -1.0809 | 3.6E-05 | ncRNA |
| SPBPB21E7.07 | aes1 | -1.0504 | 6.4E-25 | protein_coding |
| SPAC1F7.08 | fio1 | -1.0350 | 3.4E-02 | protein_coding |

**b**

| gene_ID | external_gene_ID | log2_FC | adj. p-value | gene_biotype |
|---|---|---|---|---|
| SPNCRNA.466 | SPNG2151 | 4.2720 | 2.13E-02 | ncRNA |
| SPAC977.05c | SPAC977.05c | 4.1810 | 5.59E-03 | protein_coding |
| SPBC1348.06c | SPBC1348.06c | 4.0169 | 8.64E-03 | protein_coding |
| SPMITTRNASER.01 | SPMITTRNASER.01 | 2.5719 | 1.10E-05 | tRNA |
| SPNCRNA.287 | SPNCRNA.1300,SPNG1093 | 2.5602 | 2.39E-12 | ncRNA |

| | | | | |
|---|---|---|---|---|
| SPNCRNA.100 | SPNCRNA.100 | 2.5543 | 7.41E-09 | ncRNA |
| SPMITTRNALYS.01 | SPMITTRNALYS.01 | 2.3763 | 1.32E-11 | tRNA |
| SPNCRNA.1656 | nup120-antisense-1 | 2.3445 | 1.38E-05 | ncRNA |
| SPCC1393.10 | ctr4 | 2.2131 | 3.19E-201 | protein_coding |
| SPAC1F8.04c | SPAC1F8.04c | 2.1839 | 3.42E-15 | protein_coding |
| SPBP4H10.09 | rsv1 | 1.9912 | 6.16E-04 | protein_coding |
| SPNCRNA.1282 | SPNCRNA.1282 | 1.8133 | 9.99E-11 | ncRNA |
| SPNCRNA.1677 | doa10-antisense-1 | 1.7962 | 8.50E-03 | ncRNA |
| SPMITTRNAMET.01 | SPMITTRNAMET.01 | 1.7436 | 1.54E-03 | tRNA |
| SPCC1235.14 | ght5 | 1.7393 | 8.95E-134 | protein_coding |
| SPNCRNA.601 | SPNCRNA.601 | 1.6964 | 1.15E-07 | ncRNA |
| SPAC1F8.03c | str3 | 1.6478 | 2.32E-07 | protein_coding |
| SPAC1142.05 | ctr5 | 1.6164 | 3.39E-148 | protein_coding |
| SPBPB2B2.05 | SPBPB2B2.05 | 1.5930 | 1.59E-03 | protein_coding |
| SPNCRNA.1558 | qcr10-antisense-1 | 1.4771 | 4.57E-08 | ncRNA |
| SPRRNA.01 | 21S_rRNA | 1.4280 | 7.37E-12 | rRNA |
| SPNCRNA.1307 | SPBPB10D8.02c-antisense-1 | 1.4233 | 9.50E-19 | ncRNA |
| SPNCRNA.945 | SPNCRNA.945 | 1.4199 | 5.60E-07 | ncRNA |
| SPAC1F8.06 | pfl8 | 1.3844 | 2.63E-05 | protein_coding |
| SPNCRNA.983 | tsn1-antisense-1 | 1.3631 | 6.86E-03 | ncRNA |
| SPRRNA.02 | 15S_rRNA | 1.3599 | 1.52E-10 | rRNA |
| SPBP4H10.10 | rbd3 | 1.3464 | 1.10E-09 | protein_coding |
| SPCC330.06c | pmp20 | 1.3422 | 2.45E-49 | protein_coding |
| SPBC11B10.02c | his3 | 1.3419 | 2.23E-54 | protein_coding |
| SPAC4G8.03c | puf5 | 1.2856 | 8.00E-49 | protein_coding |
| SPBC1778.04 | spo6 | 1.2527 | 2.95E-07 | protein_coding |
| SPNCRNA.634 | shk2-antisense-1 | 1.2495 | 5.43E-64 | ncRNA |
| SPNCRNA.1557 | lid2-antisense-1 | 1.2051 | 2.49E-03 | ncRNA |
| SPAC56F8.15 | SPAC56F8.15 | 1.2029 | 8.85E-28 | protein_coding |
| SPNCRNA.1438 | cdc2-antisense-1 | 1.1829 | 6.63E-03 | ncRNA |
| SPBC1289.14 | SPBC8E4.10c | 1.1804 | 2.69E-03 | protein_coding |
| SPNCRNA.1409 | SPNCRNA.1409 | 1.1604 | 6.19E-07 | ncRNA |
| SPNCRNA.33 | prl33 | 1.1499 | 3.74E-02 | ncRNA |
| SPCC1739.08c | SPCC1739.08c | 1.1300 | 2.14E-04 | protein_coding |
| SPBC1D7.02c | scr1 | 1.1176 | 2.29E-42 | protein_coding |
| SPNCRNA.1330 | SPNCRNA.1330 | 1.1154 | 4.05E-02 | ncRNA |
| SPBC23G7.08c | rga7 | 1.1060 | 7.65E-27 | protein_coding |
| SPNCRNA.892 | tif211-antisense-1 | 1.1049 | 2.85E-04 | ncRNA |
| SPNCRNA.953 | SPNCRNA.953 | 1.0861 | 1.32E-35 | ncRNA |
| SPNCRNA.1031 | wsp1-antisense-1 | 1.0573 | 1.98E-04 | ncRNA |
| SPNCRNA.875 | gcv1-antisense-1 | 1.0473 | 2.35E-08 | ncRNA |
| SPNCRNA.1378 | cbp6-antisense-1 | 1.0417 | 6.87E-09 | ncRNA |
| SPNCRNA.774 | SPNCRNA.774 | 1.0215 | 1.55E-02 | ncRNA |

| | | | | |
|---|---|---|---|---|
| SPCC794.01c | gcd1 | 1.0186 | 1.95E-09 | protein_coding |

#### **Supplementary Table 3:** *S. pombe* strains used in this study

| Strain | Genotype | Source | Figure |
|---|---|---|---|
| FP1546 | $h^-$ ade6-M216 leu1-32 ura4-D18 his3-D1 lsm4::lsm4-myc13-natMX6 pof8::kanMX6 [pDBlet-Pof8] | Páez-Moscoso et al 2018 | S1a |
| FP1547 | $h^-$ ade6-M216 leu1-32 ura4-D18 his3-D1 lsm4:: lsm4-myc13-natMX6 pof8::kanMX6 [pDBlet-3xFLAGPof8] | Páez-Moscoso et al 2018 | S1a |
| FP1913 | $h^?$ ade6-M21? leu1-32 ura4-D18 his3-D1 bmc1(SPBC2A9.10)::natMX6 [ura4, pDBlet] | This study | S4b,c |
| FP1914 | $h^?$ ade6-M21? leu1-32 ura4-D18 his3-D1 bmc1(SPBC2A9.10)::natMX6 [pDBlet–bmc1 WT] | This study | S4b,c |
| FP1915 | $h^?$ ade6-M21? leu1-32 ura4-D18 his3-D1 bmc1(SPBC2A9.10)::natMX6 [pDBlet–bmc1-Twinstrep] | This study | S4b,c |
| FP1916 | $h^?$ ade6-M21? leu1-32 ura4-D18 his3-D1 bmc1(SPBC2A9.10)::natMX6 [pDBlet–bmc1-Twinstrep F246A] | This study | S4b,c |
| FP1917 | $h^?$ ade6-M21? leu1-32 ura4-D18 his3-D1 bmc1(SPBC2A9.10)::natMX6 [pDBlet–bmc1-Twinstrep VLD39-41AAA] | This study | S4b,c |
| FP1918 | $h^?$ ade6-M21? leu1-32 ura4-D18 his3-D1 bmc1(SPBC2A9.10)::natMX6 [pDBlet–bmc1-Twinstrep GCN43-45ACA] | This study | S4b,c |
| FP1919 | $h^?$ ade6-M21? leu1-32 ura4-D18 his3-D1 bmc1(SPBC2A9.10)::natMX6 [pDBlet–bmc1-Twinstrep EPQ188-190APA] | This study | S4b,c |
| PP137 | $h^+$ ade6-M216 leu1-32 ura4-D18 his3-D1 | Lab stock | 3a, S2 |
| PP138 | $h^-$ ade6-M216 leu1-32 ura4-D18 his3-D1 | Lab stock | 3a, 3b, 3c, 3d, S2b,c,d,e, S3, S5, S6 |
| PP139 | $h^-$ ade6-M210 leu1-32 ura4-D18 his3-D1 | Lab stock | 3a, S2 |
| PP265 | $h^-$ | ATCC | S2g |
| PP1723 | $h^-$ ade6-M216 leu1-32 ura4-D18 his3-D1 pof8::kanMX6 | Páez-Moscoso et al 2018 | 3a, 3b, 3d |
| PP1797 | $h^?$ ade6-M21? leu1-32 ura4-D18 his3-D1 lsm4::lsm4-myc13-nat aur1::[pCST159-ter1] pof8::kanMX6 | This study | 5b |
| PP1839 | $h^?$ ade6-M210/M216 ura4-D18 leu1-32 his3-D1 pof8::3xFLAG-pof8-kanMX6 | This study | 3c, S5 |
| PP1843 | $h^-$ ade6-M216 leu1-32 ura4-D18 his3-D1 thc1 (SPCC18B5.09c):: thc1-TEV-2xV5-natMX6 | This study | 2a, 3c, S5 |
| PP1844 | $h^-$ ade6-M216 leu1-32 ura4-D18 his3-D1 bmc1(SPBC2A9.10)::bmc1-TEV-TwinStrep-natMX6 | This study | 2b, 3c, S5 |
| PP1845 | $h^-$ ade6-M216 ura4-D18 leu1-32 his3-D1 pof8::3xFLAG-pof8-kan thc1 (SPCC18B5.09c):: thc1-TEV-2xV5-natMX6 | This study | 2a, 4e, 4f |
| PP1846 | $h^-$ ade6-M216 ura4-D18 leu1-32 his3-D1 pof8::3xFLAG-pof8-kan bmc1(SPBC2A9.10)::bmc1-TEV-TwinStrep-natMX6 | This study | 2b, 4g, 4h |
| PP1847 | $h^-$ ade6-M216 leu1-32 ura4-D18 his3-D1 thc1 (SPCC18B5.09c)::his3 | This study | 3a, 3b, 3d, S2b,e, S3, S6 |
| PP1857 | $h^?$ ade6-M210/M216 leu1-32 ura4-D18 his3-D1 | This study | 4a, 4b |
| PP1858 | $h^?$ ade6-M210 leu1-32 ura4-D18 his3-D1 pof8::KanMX6 | This study | 4a, 4b |
| PP1859 | $h^+$ ade6-M210 leu1-32 ura4-D18 his3-D1 bmc1(SPBC2A9.10)::natMX6 | This study | 4a, 4b, S2g |
| PP1860 | $h^?$ leu1-32 ura4-D18 his3-D1 ade6-M210/ade6-M216 | This study | 3a, 3b, S2f, S3 |
| PP1861 | $h^?$ leu1-32 ura4-D18 his3-D1 ade6-M210/ade6-M216 bmc1(SPBC2A9.10)::natMX6 | This study | 3a, 3b, 3d, S2c,d,f, S3 |
| PP1862 | $h^?$ ade6-M210 leu1-32 ura4-D18 his3-D1 thc1 (SPCC18B5.09c)::his3 | This study | 4a, 4b |
| PP1863 | $h^?$ ade6-M210/216 leu1-32 ura4-D18 his3-D1 pof8::kanMX6 bmc1(SPBC2A9.10)::natMX6 | This study | 4a, 4b |
| PP1864 | $h^?$ ade6-M216 leu1-32 ura4-D18 his3-D1 pof8::kanMX6 thc1 (SPCC18B5.09c)::his3 | This study | 4a, 4b |
| PP1865 | $h^?$ ade6-M210/216 leu1-32 ura4-D18 his3-D1 bmc1(SPBC2A9.10)::natMX6 thc1 (SPCC18B5.09c)::his3 | This study | 4a, 4b |

| | | | |
|---|---|---|---|
| PP1866 | $h^?$ ade6-M210 leu1-32 ura4-D18 his3-D1 pof8::kanMX6 bmc1(SPBC2A9.10)::natMX6 thc1 (SPCC18B5.09c)::his3 | This study | 4a, 4b |
| PP1882 | $h^?$ ade6-M210 leu1-32 ura4-D18 his3-D1 smb1:: smb1-cMyc-natMX6, pof8::kanMX6 | This study | 5f, 5g |
| PP1883 | $h^?$ ade6-M210 leu1-32 ura4-D18 his3-D1 smb1:: smb1-cMyc-natMX6, thc1 (SPCC18B5.09c)::his3 | This study | 5f, 5g |
| PP1884 | $h^?$ ade6-M216 leu1-32 ura4-D18 his3-D1 smb1:: smb1-cMyc-natMX6, bmc1(SPBC2A9.10)::natMX6 | This study | 5f, 5g |
| PP1885 | $h^?$ ade6-M210/M216 leu1-32 ura4-D18 his3-D1 smb1::smb1-cMyc-natMX6 | This study | 5f, 5g |
| PP1886 | $h^?$ ade6-M216 leu1-32 ura4-D18 his3-D1 lsm4::lsm4-cMyc-natMX6, pof8::kanMX6 | This study | 5c, 5d, 5e |
| PP1887 | $h^?$ ade6-M210 leu1-32 ura4-D18 his3-D1 lsm4::lsm4-cMyc-natMX6, thc1 (SPCC18B5.09c)::his3 | This study | 5c, 5d, 5e |
| PP1888 | $h^?$ ade6-M216 leu1-32 ura4-D18 his3-D1 lsm4::lsm4-cMyc-natMX6, bmc1(SPBC2A9.10)::natMX6 | This study | 5c, 5d, 5e |
| PP1889 | $h^?$ ade6-M210/M216 leu1-32 ura4-D18 his3-D1 lsm4::lsm4-cMyc-natMX6 | This study | 5c, 5d, 5e |
| PP1892 | $h^?$ ade6-M216 leu1-32 ura4-D18 his3-D1 thc1 (SPCC18B5.09c)::thc1-TEV-2xV5-natMX6 bmc1(SPBC2A9.10)::bmc1-TEV-Twinstrep-natMX6 | This study | 2c, 4c, 4d |
| PP1894 | $h^?$ ade6-M216 leu1-32 ura4-D18 his3-D1 bmc1(SPBC2A9.10)::bmc1-TEV-Twinstrep-natMX6 | This study | 2c |
| PP1895 | $h^?$ ade6-M216 leu1-32 ura4-D18 his3-D1 pof8::kanMX6 thc1 (SPCC18B5.09c)::thc1-TEV-2xV5-natMX6 bmc1(SPBC2A9.10)::bmc1-TEV-Twinstrep-natMX6 | This study | 4c, 4d |
| PP2014 | $h^?$ ade6-M21? leu1-32 his3-D1 ura4-D18 pof8::3xFLAG-pof8-kanMX6 | This study | 6a, 6b |
| PP2015 | $h^?$ ade6-M21? leu1-32 his3-D1 ura4-D18 pof8::3xFLAG-pof8-kanMX6 thc1 (SPCC18B5.09c)::his3 | This study | 6a, 6b |
| PP2016 | $h^?$ ade6-M21? leu1-32 his3-D1 ura4-D18 pof8::3xFLAG-pof8-kanMX6 bmc1(SPBC2A9.10)::natMX6 | This study | 6a, 6b |
| PP2017 | $h^?$ ade6-M21? leu1-32 his3-D1 ura4-D18 pof8::3xFLAG-pof8-kanMX6 thc1 (SPCC18B5.09c)::his3 bmc1(SPBC2A9.10)::natMX6 | This study | 6a, 6b |
| PP2024 | $h^?$ ade6-M21? ura4-D18 leu1-32 his3-D1 thc1::thc1-TEV-2xV5-natMX6 lsm4:: lsm4-cMyc-natMX6 | This study | 6c, 6e, 6g |
| PP2025 | $h^?$ ade6-M216 ura4-D18 leu1-32 his3-D1 bmc1::bmc1-TEV-Twinstrep-natMX6 lsm4:: lsm4-cMyc-natMX6 | This study | 6d, 6f, 6g |
| PP2038 | $h^?$ ade6-M21? ura4-D18 leu1-32 his3-D1 3xFLAG-pof8-kanMX6 thc1 (SPCC18B5.09c):: thc1-TEV-2xV5-natMX6 bmc1(SPBC2A9.10)::natMX6 | This study | 4e, 4f |
| PP2039 | $h^?$ ade6-M21? ura4-D18 leu1-32 his3-D1 3xFLAG-pof8-kanMX6 bmc1(SPBC2A9.10)::bmc1-TEV-TwinStrep-natMX6 thc1(SPCC18B5.09c)::his3 | This study | 4g, 4h |

**Supplementary Table 4:** Oligonucleotides and gene synthesis products used to generate deletion, fusion and integration constructs

| Product description | Primer # | Sequence |
|---|---|---|
| *Nat MX6* for *bmc1* deletion | BLoli6066/BLoli2491 | 5'-TTTAGCTTGCCTCGTCCCCG-3'/5'-TGGATGGCGGCGTTAGTATC-3' |
| *bmc1* 5' homology | BLoli7736/BLoli7737 | 5'-ATCCTGAAGCGATGATGCCA-3'/ 5'-CGGGGACGAGGCAAGCTAAACCCAAGTCGAGGAGGTTTTT-3' |
| *bmc1* 3' homology | BLoli7738/BLoli7739 | 5'-GATACTAACGCCGCCATCCATTGTCTAGTAAAACGTTAAG-3'/5'-TTGGCGAGTATAACCAATGT-3' |
| *bmc1*::natMX6 | BLoli7736/BLoli7739 | 5'-ATCCTGAAGCGATGATGCCA-3'/5'-TTGGCGAGTATAACCAATGT-3' |
| his3$^{+}$ for *thc1* deletion | BLoli7486/BLoli7487 | 5'-GTTTTGAAGACGGTGATACACGTTGTAATG-3'/5'-ATTTATCTGTTTGCTTATCGAACTATACGG-3' |
| *thc1* 5' homology | BLoli7484/BLoli7485 | 5'-CAATAATAACTTTGCTTACGATTAATAG-3'/5'-TGTATCACCGTCTTCAAAACTTTTGGTACC-3' |
| *thc1* 3' homology | BLoli7488/BLoli7489 | 5'CGATAAGCAAACAGATAAATTAGAACACAGC-3'/5'-GAGACTAATTGGGTTAAACAAAG-3' |
| *thc1*::his3 | BLoli7484/BLoli7489 | 5'-CAATAATAACTTTGCTTACGATTAATAG-3'/5'-GAGACTAATTGGGTTAAACAAAG-3' |
| *bmc1* 5' homology for tagging | BLoli7718/BLoli7709 | 5'-AGCTCTCGAATTGGCCCTG-3'/5'-TTTAGAAGTGTTATTTCTCAAATTGAGGATGACTCCATG-3' |
| *bmc1* 3' homology for tagging | BLoli7541/BLoli7542 | 5'-GCCATCCAGTTAATTGTCTAGTAAAACGTTAAGAATAG/5'-TGATGTTGGCGAGTATAAC-3' |
| natMX6 cassette for *bmc1* tagging | BLoli7710/BLoli7540 | 5'-TGAGAAATAACACTTCTAAATAAGCGAATTTC-3'/5'-TAGACAATTAACTGGATGGCGGCGTTAG-3' |
| *bmc1*-Twinstrep::natMX6 | BLoli7718/BLoli7542 | 5'-AGCTCTCGAATTGGCCCTG-3'/5'-TGATGTTGGCGAGTATAAC-3' |
| *thc1* 5' homology for tagging | BLoli7711/BLoli7712 | 5'-TCTTAAGATATTTGGGCTATAAATG-3'/5'-TTTAGAAGTGTTATTACGTGGAATCTAATCC-3' |
| *thc1* 3' homology for tagging | BLoli7715/BLoli7716 | 5'-GCCATCCAGTACAGATAAATTAGAACACAGC-3'/5'-AACAAAGTAGTAACCAAGG-3' |
| natMX6 cassette for *thc1* tagging | BLoli7713/BLoli7714 | 5'-CACGTAATAACACTTCTAAATAAGCGAATTTCTTATGATTTATG-3'/5'-ATTTATCTGTACTGGATGGCGGCGTTAG-3' |
| Thc1-2xV5::natMX6 | BLoli7711/BLoli7716 | 5'-TCTTAAGATATTTGGGCTATAAATG-3'/5'-AACAAAGTAGTAACCAAGG-3' |
| 3xFLAG-Pof8 5' homology for tagging | BLoli6676/BLoli6400 | 5'-AAAAGAATTCAACATGGCAACTGCGACCAA-3'/5'-TTTAGAAGTGTTACTTTTTAACATACGCCAATAATTC-3' |
| Pof8-kanMX6 3' homology for tagging | BLoli6401/BLoli6141 | 5'-GGCGTATGTTAAAAAAGTAACACTTCTAAATAAGCG-3'/5'-GCTTTCTTATTTGTAGAGACAATTG-3' |
| 3xFLAG-Pof8-kanMX6 | BLoli6676/BLoli6141 | 5'-AAAAGAATTCAACATGGCAACTGCGACCAA-3'/5'-GCTTTCTTATTTGTAGAGACAATTG-3' |
| Cloning of bmc1-Twinstrep into pDBlet | BLoli8069/BLoli8070 | 5'-CCGATAAGCTTAAACTATCTTAACCTGTCTACG-3'/5'-GGTGGCGGCCGCTACAGTTTTGGTATACCAGG-3' |
| bmc1 F246A 5' arm | Bloli8069/BLoli8006 | 5'-CCGATAAGCTTAAACTATCTTAACCTGTCTACG-3'/5'-GTTCGTTTGGCAGCATTCTTGTAC-3' |
| bmc1 F246A 3' arm | BLoli8007/BLoli8070 | 5'-GTACAAGAATGCTGCCAAACGAAC-3'/5'-GGTGGCGGCCGCTACAGTTTTGGTATACCAGG-3' |
| bmc1 VLD39–41AAA 5' arm | Bloli8069/BLoli7792 | 5'-CCGATAAGCTTAAACTATCTTAACCTGTCTACG-3'/5'-CATTATTGCATCCTATCGCTGCGGCTGAAGCCTC-3' |
| bmc1 VLD39–41AAA 3' arm | BLoli7793/BLoli8070 | 5'-CAGCCGCAGCGATAGGATGCAATAATGGGAC-3'/5'-GGTGGCGGCCGCTACAGTTTTGGTATACCAGG-3' |
| bmc1 GCN43–45ACA 5' arm | Bloli8069/BLoli7795 | 5'-CCGATAAGCTTAAACTATCTTAACCTGTCTACG-3'/5'-GAGCAGACACTGTCCCATTAGCGCACGCTATTGCT-3' |
| bmc1 GCN43–45ACA 3' arm | BLoli7796/BLoli8070 | 5'-GTGCGCTAATGGGACAGTGTCTGCTCAAATTG-3'/5'-GGTGGCGGCCGCTACAGTTTTGGTATACCAGG-3' |
| bmc1 EPQ188–190APA 5' arm | Bloli8069/BLoli7802 | 5'-CCGATAAGCTTAAACTATCTTAACCTGTCTACG-3'/5'-TCAAGTACGAGTCCCATCCGGCAGGTGCTAAAATAAG-3' |
| bmc1 EPQ188–190APA 3' arm | BLoli7803/BLoli8070 | 5'-ACCTGCCGGATGGGACTCGTACTTGAAAGCTG-3'/5'-GGTGGCGGCCGCTACAGTTTTGGTATACCAGG-3' |
| Bmp1 full length PCR verification | BLoli8048/BLoli7740 | 5'-GAGAGACTAGCACCAATGTATCC-3'/5'-TAGCTCTTGAAAAGGTACGG-3' |

| | | |
|---|---|---|
| 5' arm bmp1::natMX6 PCR verification | BLoli7735/BLoli3688 | 5'-CACTACAATTCAAGACTACC-3'/5'-GTAAGCCGTGTCGTCAAGAG-3' |
| 3' arm bmp1::natMX6 PCR verification | BLoli7740/BLoli3791 | 5'-TAGCTCTTGAAAAGGTACGG-3'/5'-GGCGCTCTACATGAGCATGC-3' |
| 5' arm bmp1 PCR verification | BLoli8048/BLoli8049 | 5'-GAGAGACTAGCACCAATGTATCC-3'/5'-AGGGAATCAGGCAAACATTTAAG-3' |
| thc1::his3 full length PCR verification | BLoli7484/BLoli7717 | 5'-CAATAATAACTTTGCTTACGATTAATAG-3'/5'-TGAAGCACCAAGATACGAGT-3' |

#### Gene synthesis products

| | | |
|---|---|---|
| thc1-TEV-2xV5 fragment | BLgs1466 | 5'-<br>TGCTGTATCGGAGCCTAATTCTAGTGGTCTTAAGATATTTGGGCTATAAATGATTGAAAATTGGC<br>AATCTTGTTCACGCTGATCAACAATAAAAAAACGGCTACTTAGTATTTACCCTATACTCTTAGCA<br>TGGATACCTTCAATATTCATTGAAGTTGTAGTTTTAGCTTATTTGGTACCAAAAGTTTTGAAGAAT<br>GGAAGAGAAAAATACTGTTTCTTTATCTAAGCATATTGAACGTCCAGTAGAAGTTGTTGAAAGTC<br>ATTCTACGTACATTTTAAGTGCACAAGGACTTTATCTTACAGAACGCGTTTTAAGAAGTTATTTTA<br>AACAACCTGATCTAATTATTACTTGGAAGGATAGTATGAGAGCTTACCTGACATTTTCTTCGCCG<br>CAGGAAGCTCAAAAGGCTTACTTAGATTCACTTCGTTGGGGCAGTCAACTGAATGCTATCATTAA<br>ACCATTCTACGGTTCGCACGATGAGGTACTTCGTTTATGTAAAAGGAAAAGAATTATTCCACTAC<br>AAAATTTCTTGACTTCAGGTCTCGAGCCAACCACTGAGGATCTGTACTTTCAGAGCGATAACGAT<br>GGTAAGCCTATCCCTAACCCTCTCCTCGGTCTCGATTCTACGGGCGGAGGCTCCGGCGGCGGCTC<br>TGGAGGATCTGCAGGTAAGCCAATACCCAACCCACTTCTTGGATTAGATTCCACGTAATAACACT<br>TCTAAATAAGCGAAT-3' |
| bmc1-TEV-Twin-Streptag fragment | BLgs1465 | 5'-<br>AGCTCTCGAATTGGCCCTGTACGCAATCCTGGTTCCATTGTAGAAGACCAGTTTAATTATTACCCC<br>ATTTCAAGCATTAAAAAGTTTTCCAGGATACCAGTGCAACTTCAACCACCTCTCAATAAGCAAAA<br>TTTCCCTCACAATATAGAATTTGAGACCGCTGACTTCTTGCGCTGGGAATCGAAACGAAAATTCA<br>AAATAATACTAGCATTATCCGTATCTAAATGGGTGCATCTAAATAACCACGATGAAGGAATCATT<br>AAATTCTTTGGGAAGATTAGTTCTTTATTGGAAACGAATGGTGTTCTTATTTTAGAACCTCAAGGA<br>TGGGACTCGTACTTGAAAGCTGCAAAAAAAATATCTGTAAGTCACGACTATCAATACTTCTAACT<br>CTTACTTTTCTAGGTTTTTAATCAAACACCTGAGAACCTCAAAATCCAACCTGATGCGTTTGAACA<br>TTTGCTTAATCAAGCAGGACTAGTGCTTGAATACAGTATCGAACCTCAAGTAAATAACTCTGAGT<br>ACAAGAATTTTGCCAAACGAACAATGTATATCTATAAAAAAAAAGGAATTGGAATCATAAAACT<br>ATTAACTTCTACTCTCGAGCCAACAACTGAAGATTTATATTTTCAATCTGACAATGATTGGTCTCA<br>CCCACAGTTTGAAAAAGGCGGAGGCTCCGGCGGCGGCTCTGGAGGATCTGCATGGAGTCATCCT<br>CAATTTGAGAAATAACACTTCTAAATAAGCGAATTTC-3' |

**Supplementary Table 5:** Oligonucleotides used for RT-qPCR

| Product | Primer # | Sequence | Source |
|---|---|---|---|
| TER1 all | BLoli7827 | 5'-CAGTGTACGTGAGTCTTCTGCCTT-3' | Ref. 15 |
| | BLoli7828 | 5'-CAAAAATTCGTTGTGATCTGACAAGC-3' | Ref. 15 |
| TER1 exon 2 | BLoli7825 | 5'-AATTGCGTATTTAGTAAGAACGCG-3' | Ref. 16 |
| | BLoli7826 | 5'-GATTCATCACTTTCTCAAAATTTTGAAACCG-3' | Ref. 16 |
| act1 | BLoli7829 | 5'-GGATTCCTACGTTGGTGATGA-3' | Ref. 16 |
| | BLoli7830 | 5'-CGTTGTAGAAAGTGTGATGCC-3' | Ref. 16 |
| his1 | BLoli7818 | 5'-CGAAGACGTGCTTCAGCGA-3' | Ref. 15 |
| | BLoli7819 | 5'-TGTCCACCTCGGAATCACTG-3' | Ref. 15 |
| snR101 | BLoli7866 | 5'-CGCTCTAGAAATTGGAATGAG-3' | This study |
| | BLoli7867 | 5'-TCTTAAAGGTGTGTCTCTCC-3' | This study |
